# Computational drug repurposing identified Artemisinin and Mebendazole as potential inhibitors of virulence-associated proteins SKSR and essential kinases CpCDPK1 of *Cryptosporidium parvum*

**DOI:** 10.64898/2026.05.17.725751

**Authors:** Parveen, Deeksha Saini, Munendra Kumar, Kapinder, Anusha Singh, Nida Jamil Khan, Nikhat Manzoor, Manish Sharma, Prateek Kumar

## Abstract

*Cryptosporidium parvum* is a protozoan parasite responsible for cryptosporidiosis, significantly threatening immunocompromised individuals, particularly HIV/AIDS patients, by causing severe diarrhea and potential mortality. Current treatments are largely ineffective, prompting investigations into new therapeutic options. This study evaluated two antiparasitic drugs: Mebendazole, used for helminth infections, and Artemisinin, used for malaria. The SKSR gene family encodes virulence factors in *C. parvum*, and Calcium-dependent protein kinase1 (CpCDPK1) regulates the life cycle of *C. parvum;* targeting these proteins may reduce growth and infection in hosts. In the current study, molecular docking was conducted taking Mebendazole and Artemisinin drugs as ligands, SKSR gene family and CpCDPK1 proteins as drug targets. Results with SKSR showed binding energy of -4.9 kcal/mol, -6.72 kcal/mol for Mebendazole and Artemisinin, respectively. Whereas, with CpCDPK1, the binding energies were -6.44 kcal/mol, -9.18 kcal/mol for Mebendazole and Artemisinin, respectively. Docking of Nitazoxanide (an in-use drug for *C. parvum*) with SKSR and CpCDPK1 revealed binding energies -4.2 kcal/mol, -4.81 kcal/mol, respectively. The stability of the proteins (targets) upon binding to the ligands was assessed by performing all-atom MD simulations for 100ns using the GROMACS package. No major variations were observed upon binding of Artemisinin and Mebendazole to SKSR and CpCDPK1. The findings of MD simulations imply that both proteins maintain their stability upon binding of Artemisinin and Mebendazole. Molecular Docking and MD simulation studies suggest that Artemisinin and Mebendazole are potential candidates for repurposing in the treatment of *C. parvum* infections, with recommendations for *in vitro* studies to validate these findings.

## 1. Introduction

*Cryptosporidium parvum* infection leads to diarrhea, which ultimately results in malnutrition and finally death (Katiyar et al. 2022; He et al. 2024; Al-Mamun et al. 2024). Other symptoms are abdominal pain and acute gastroenteritis. This disease condition is known as Cryptosporidiosis, which is a major cause behind the food and water-borne diseases in both developing and developed countries (Katiyar et al. 2022; Wang et al. 2023). This disorder is found to be severe in immunocompromised and young children, which ultimately leads to mortality. Upon literature studies, it was found that the virulence mechanism of *Cryptosporidium* is not well-understood (He et al. 2024). The hosts of this pathogen are several vertebrates and human beings. In humans, more than 20 species of *Cryptosporidium* have been identified. More than 90% cases of *Cryptosporidiosis* are due to *Cryptosporidium parvum* and *Cryptosporidium hominis*. In case of humans, the main cause of disease development is *Cryptosporidium parvum* (Gerace et al. 2019). Infection with this pathogen is very dominant in the case of immunocompromised individuals, such as patients suffering from HIV/AIDS. In these *Cryptosporidium parvum* patients, females shed the oocytes in the stool during the entire duration, and severe watery diarrhea takes place for more than two months; these symptoms ultimately lead to the death of the patient. Several other symptoms, such as anorexia, fatigue, cramps, vomiting, low fever, and nausea, also exist during the infection. There are some antiretroviral therapies (ART) to boost the immune system of HIV patients, due to which infection of *Cryptosporidium parvum* can be controlled to some extent. Due to a lack of good therapeutic options, the co-infection of *Cryptosporidium parvum* in HIV infected patients is a big threat for such patients (Wang et al. 2018). There is an urgent need to explore good therapeutic options to combat the co-infection of *Cryptosporidium parvum* in HIV infected/immunocompromised patients (Wang et al. 2018). The current *in silico* study was planned to explore new therapeutic options to combat *Cryptosporidium parvum*. In the present investigation, to address technical and financial barriers of drug development to combat cryptosporidiosis, we planned a drug repurposing screen of Mebendazole and Artemisinin drugs against *Cryptosporidium parvum*. Mebendazole and Artemisinin are currently in use to treat helminth infections and malaria, respectively. The goal of our study is to evaluate the inhibitory effect of Mebendazole and Artemisinin against the apicomplexan protozoan parasite *Cryptosporidium parvum*. The SKSR gene family is identified to code proteins which acts as key virulence factors in *Cryptosporidium parvum*. Calcium-dependent protein kinase1 (CpCDPK1) which is responsible for regulating the life cycle of *Cryptosporidium parvum*. It acts as a key regulator in intracellular growth, invasion of host cells, and control of the gliding movements of the parasite. Inhibition of SKSR gene products and CpCDPK1 by drugs Mebendazole and Artemisinin may lead to inhibiting the growth of *C. parvum* and finally lead to controlling the Cryptosporidiosis (He et al. 2024; Bouzid et al. 2013). During the present investigation, molecular docking and MD simulation results revealed that both proteins (i.e., SKSR and CpCDPK1) maintain their stability upon binding of Artemisinin and Mebendazole. Hence, it could be inferred that the proteins retained their respective conformations and remained stable during the course of MD simulations. In conclusion, Molecular Docking and MD simulation studies indicated that Artemisinin and Mebendazole can be repurposed to treat *Cryptosporidium parvum* infections. We further recommend the *in vitro* studies for the validation of these *in silico* study findings.

## 2. Materials and Methods

### 2.1 Mebendazole and Artemisinin drugs as ligand

Drugs Mebendazole (PubChem CID 4030) and Artemisinin (PubChem CID 68827), which are currently used to treat helminthic and *Plasmodium falciparum* infections, respectively, were downloaded from PubChem and were used as ligands. Currently approved drug Nitazoxanide (PubChem CID 41684) to treat *Cryptosporidium parvum* was also taken as ligand to compare the docking results. The smiles of proposed repurposing drugs Mebendazole, Artemisinin, and in-use drug (Nitazoxanide) extracted from PubChem were also used for drug-likeness property analyses using ADMET Lab 3.0 (https://admetlab3.scbdd.com).

### 2.2 Drug targets of *Cryptosporidium parvum*

Targets, secreted protein of *Cryptosporidium parvum*-specific SKSR gene family (SKSR) was retrieved from AlphaFold Protein structure database, and Calcium-dependent protein kinase (CpCDPK1) was retrieved from PDB (PDB ID 3MWU). First target, secreted protein of *Cryptosporidium parvum*-specific SKSR gene family (SKSR). SKSR is responsible for the virulence of *Cryptosporidium parvum*. Second target, Calcium-dependent protein kinase 1 from *Cryptosporidium parvum* (CpCDPK1). It is responsible for regulating the life cycle of *Cryptosporidium parvum*. It acts as a key regulator in intracellular growth, invasion of host cells, and control of the gliding movements of the parasite. Inhibition of SKSR gene products and CpCDPK1 may lead to inhibiting the growth of *C. parvum* and finally lead to controlling the Cryptosporidiosis.

### 2.3 Molecular docking studies

The inhibitory effect of ligands Mebendazole, Artemisinin, and Nitazoxanide (recommended drug) against the key targets of *Cryptosporidium parvum* SKSR protein and CpCDPK1 was studied using Autodock tools. The structures of ligands were retrieved from PubChem, and structures of drug targets were retrieved from AlphaFold protein structure database and PubChem. Further, the docking results were analyzed using Protein-Ligand Interaction Profiler (PLIP) (https://plip-tool.biotec.tu-dresden.de/plip-web/plip/index) and PyMOL.

### 2.4 Molecular Dynamics (MD) Simulations

To examine the dynamics and stability of the docked complexes, all-atom molecular dynamics simulations were performed using the GROningen MAchine for Chemical Simulations (GROMACS) package (Abraham et al. 2015). Unbound proteins SKSR and CpCDPK1, along with their complexes with Artemisinin and Mebendazole, were subjected to MD simulations for 100ns. The topologies of the proteins and the ligands were prepared using Charmm36 all-atom force field (Huang et al. 2013) and the CGenFF server, respectively. All the systems were placed at 1nm from the edge of a dodecahedron box and solvated using the TIP3P water model. The salt concentration of the systems was simulated by the addition of Na and Cl ions using the gmx genion module. Subsequently, an energy minimization step was performed using the steepest decent algorithm with 50,000 steps. The apo-proteins, along with their complexes, were equilibrated using NVT and NPT ensembles for 100ps. Pressure and temperature were kept at 1 bar and 300K, respectively. The trajectory obtained at the end of the simulations was analyzed for RMSF, RMSD, Rg, and SASA using GROMACS tools, including gmx rmsf, gmx rms, gmx sasa, and gmx gyrate, respectively. Qt Grace was used for the generation and visualization of plots.

## 3. Results and Discussion

### 3.1 Drug likeness of Mebendazole and Artemisinin

An *in-silico* tool was used to identify the drug-likeness properties of proposed repurposing drugs, Mebendazole & Artemisinin and compared with the recommended drug (Nitazoxanide) for the treatment of *Cryptosporidium parvum*. The molecular weight of drug molecules plays an important role in drug likeness; according to the Lipinski rule of five, the molecular weight of drugs should be less than 500 Da. Both Mebendazole & Artemisinin have a lower molecular weight than the recommended drug Nitazoxanide. It is well recommended that the smaller the molecular weight, the higher the chances of absorption of drugs and the higher the chances of crossing the blood-brain barrier. As a result, both compounds have better BBB permeability and an effective Topological polar surface area (TPSA). According to the literature, TPSA is recommended to be less than 140 Å^2^ for potential drug molecules. Mebendazole & Artemisinin have 84.08 and 53.99 Å^2^ TPSA, respectively, which is far better than Nitazoxanide (111.43 Å^2^). In respect of Hydrogen bond acceptors (nHA) and hydrogen bond donors (nHD) of drug compounds, it should lie in the limit of nHA ≤10 and nHD ≤5, both compounds are in the recommended limit. All compounds (Mebendazole, Artemisinin, and Nitazoxanide) have zero violation of Lipinski rules of five (Table 1).

**Table 1:** Drug likeliness properties of repurposing drugs (Mebendazole and Artemisinin) and recommended drug (Nitazoxanide) for *Cryptosporidium parvum*.

| Compounds | MW (Da) | BBB | TPSA | Dense | nHA | nHD | Lipinski |
| --- | --- | --- | --- | --- | --- | --- | --- |
| Mebendazole | 295.1 | 0.284531 | 84.08 | 0.999473 | 6 | 2 | 0 |
| Artemisinin | 282.15 | 0.012941 | 53.99 | 1.025683 | 5 | 0 | 0 |
| Nitazoxanide | 307.03 | 0.978342 | 111.43 | 1.123195 | 8 | 1 | 0 |

### 3.2 Molecular Docking Studies

Docking studies revealed that Mebendazole and Artemisinin exhibit strong binding affinities for the SKSR protein responsible for the virulence of *Cryptosporidium parvum*. Results showed binding energy of -4.9 kcal/mol, -6.72 kcal/mol for Mebendazole and Artemisinin, respectively (Tables 1 & 2). The SKSR gene family is identified to code proteins that act as key virulence factors in *Cryptosporidium parvum* (He et al. 2025). By inhibiting these virulence proteins (i.e., SKSR), we can inhibit the growth and infection of this pathogen in its host. Docking results suggested that Artemisinin can inhibit the growth of *Cryptosporidium parvum* more effectively by inhibiting the SKSR gene, which can ultimately result in the death of this pathogen (Fig. 1,3). During docking studies of Mebendazole and Artemisinin with CpCDPK1, results revealed binding energy of -6.44 kcal/mol, -9.18 kcal/mol for Mebendazole and Artemisinin, respectively (Tables 2 & 3). In *Cryptosporidium parvum*, CpCPDK1 is an important signaling enzyme that regulates host cell invasion, motility of the parasite, and egress. Also, it functions to control the signaling of calcium and facilitate the discharge of organelles required for causing infection (Su et al. 2022). So, it is vital for parasites’ survival. By inhibiting this key drug target, we can inhibit the growth and infection of this pathogen. During docking studies of Nitazoxanide with SKSR and CpCDPK1, results revealed binding energies -4.2 kcal/mol, -4.81 kcal/mol, respectively (Fig. 5, 6). Our molecular docking results suggested that Mebendazole and Artemisinin can inhibit the growth and infection of *Cryptosporidium parvum* more effectively than Nitazoxanide (recommended drug) by inhibiting the CpCDPK1 protein, which can ultimately result in the death of this pathogen (Fig. 2,4). We further recommend the *in vitro* studies for the validation of these *in silico* study findings.

**Table 2:** Molecular docking studies of Mebendazole with key targets SKSR protein and CpCDPK1 of *Cryptosporidium parvum*.

| S.No. | Ligand docked with receptor complex | Binding energy (kcal/mol) | Interacting amino acids |
| --- | --- | --- | --- |
| 1. | Mebendazole (Ligand) + SKSR protein (Drug target) | -4.9 | 863A VAL, 870A VAL, 872A TYR, 872A TYR, 873A LEU, 876A PHE |
| 2. | Mebendazole (Ligand) + CpCDPK1 (Drug Target) | -6.44 | 100A GLN, 101A GLU, 153A GLU, 156A THR, 378A THR, 380A ASP, 380A ASP |

**Fig. 1:**
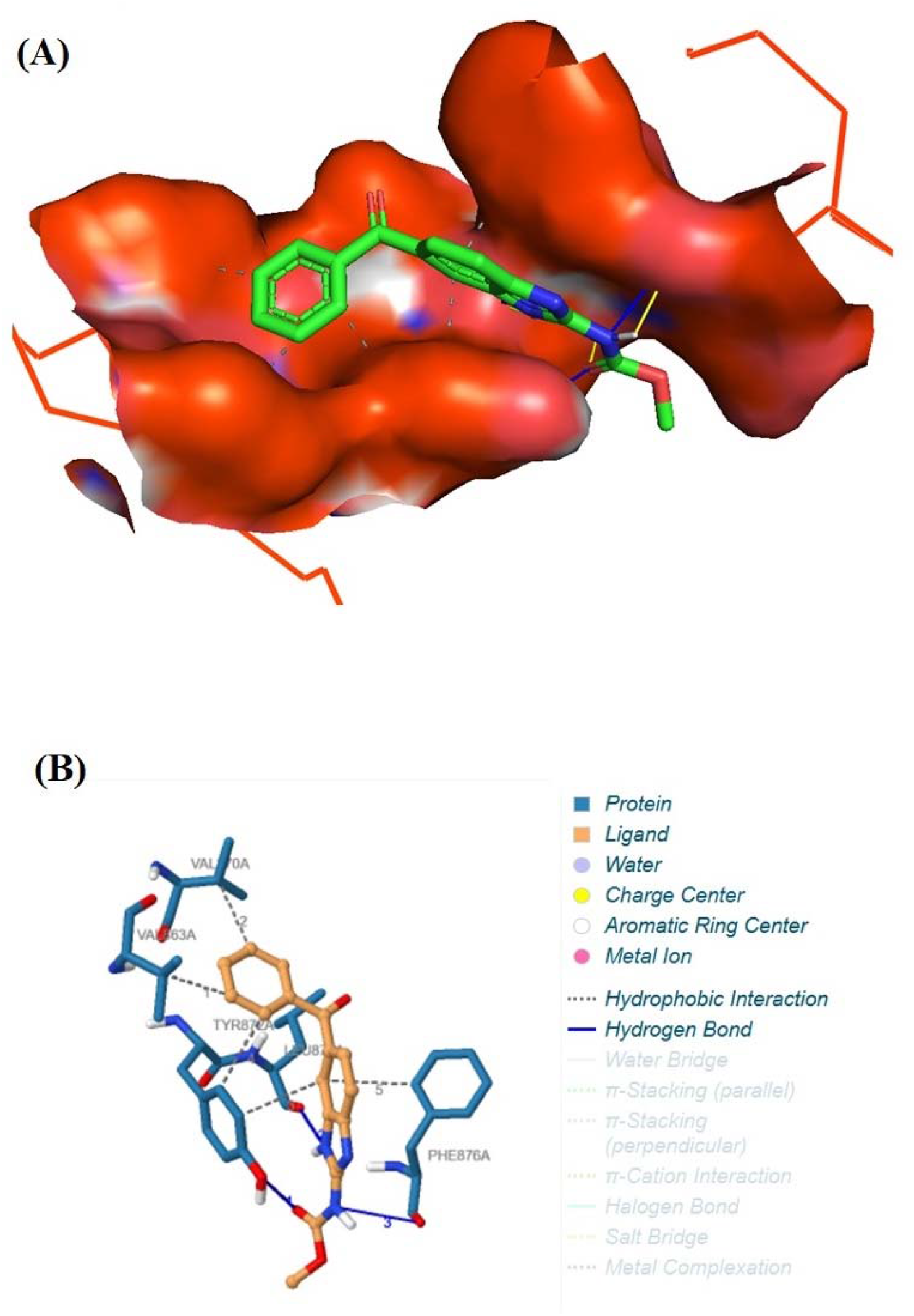
**(A.)** Enlarged view of cavity representation of pocket of SKSR protein of *Cryptosporidium parvum* where drug Mebendazole binds; **(B.)** Mebendazole-SKSR protein interaction

**Fig. 2:**
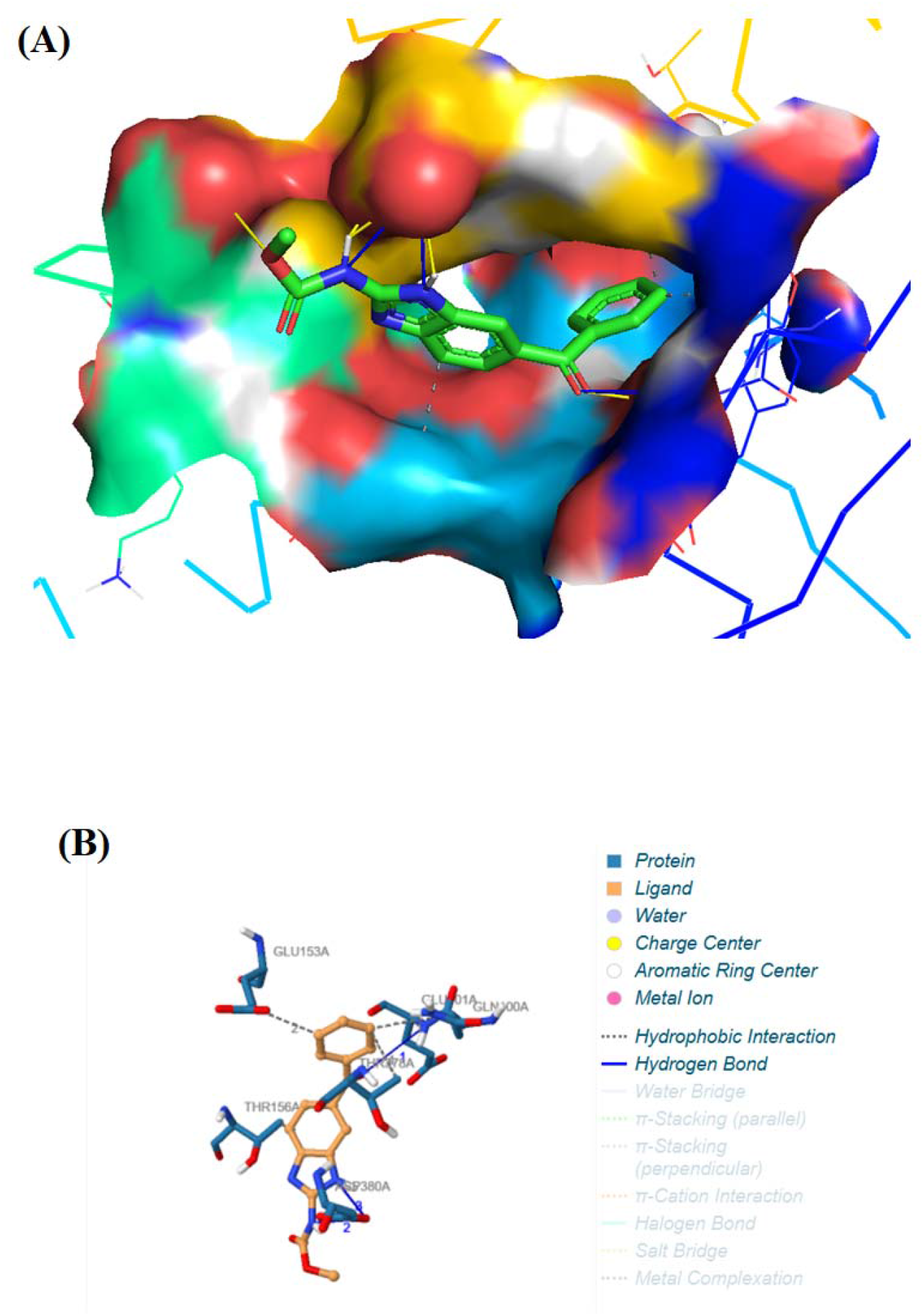
**(A.)** Enlarged view of cavity representation of pocket of key target CpCDPK1 of *Cryptosporidium parvum* where drug Mebendazole binds; **(B.)** Mebendazole-CpCDPK1 protein interaction

**Fig. 3:**
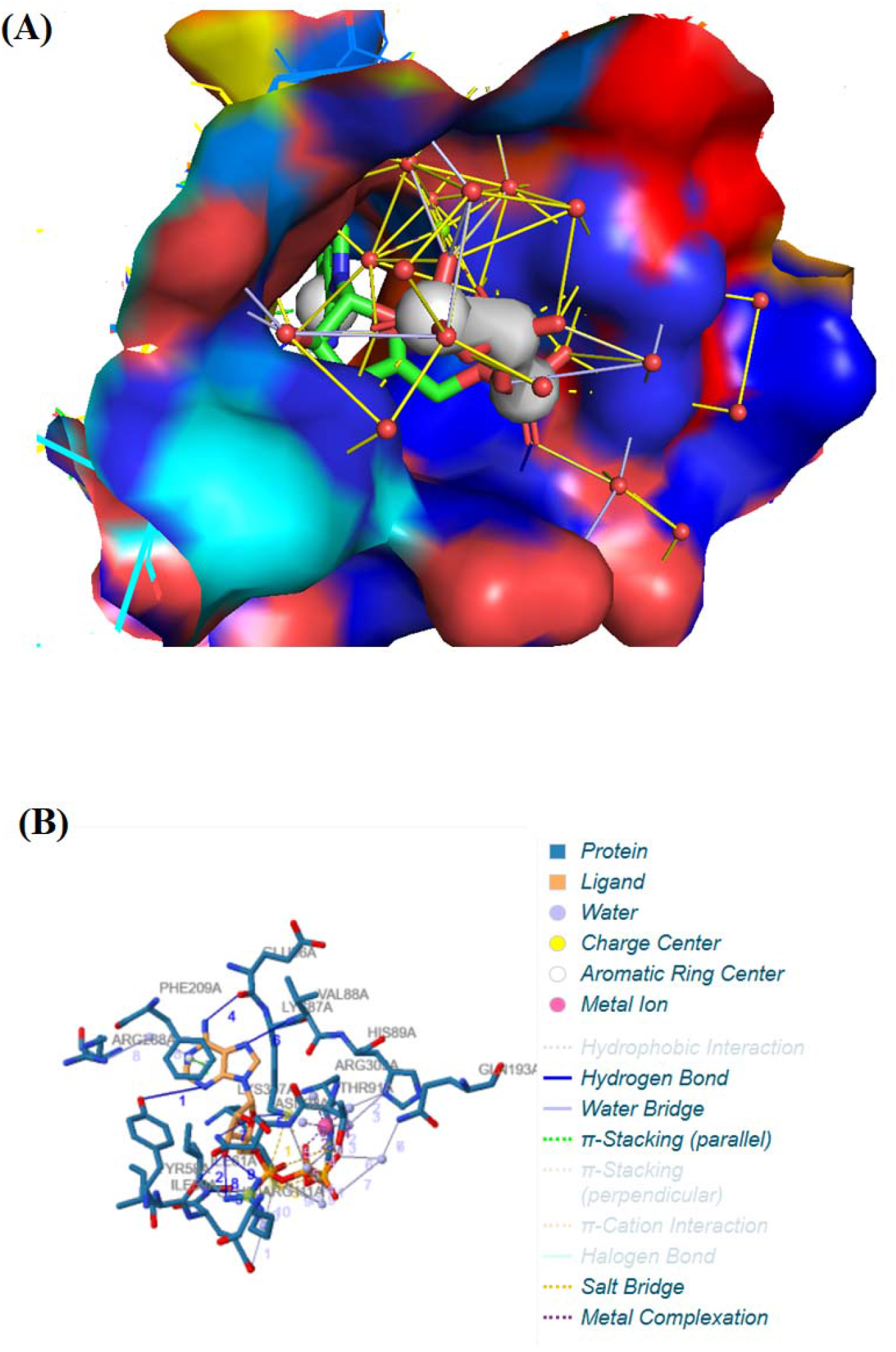
**(A.)** Enlarged view of cavity representation of pocket of key target SKSR protein of *Cryptosporidium parvum* where drug Artemisinin binds; **(B.)** Artemisinin-SKSR protein interaction

**Fig. 4:**
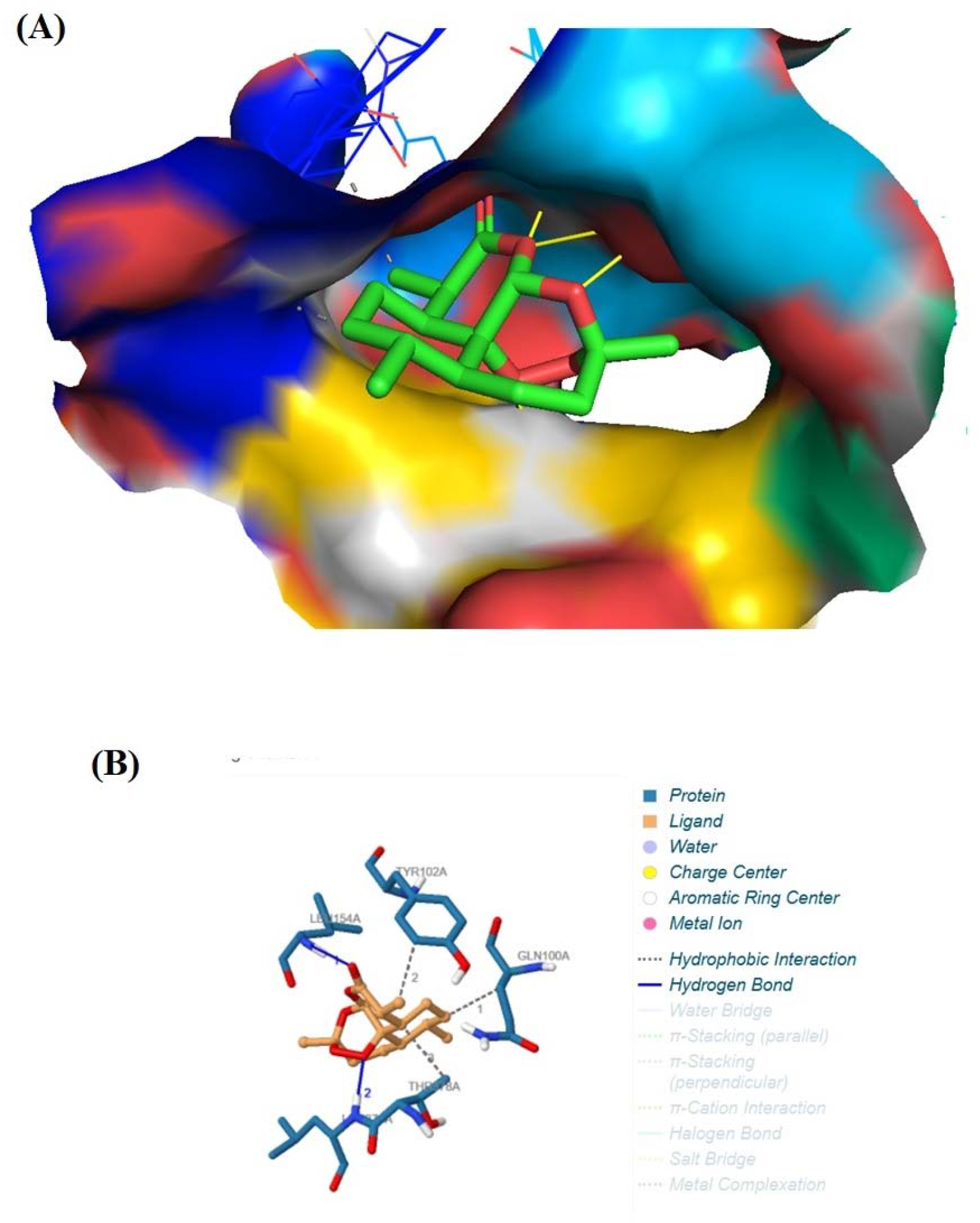
**(A.)** Enlarged view of cavity representation of pocket of key target CpCDPK1 of *Cryptosporidium parvum* where drug Artemisinin binds; **(B.)** Artemisinin-CpCDPK1 protein interaction.

**Fig. 5:**
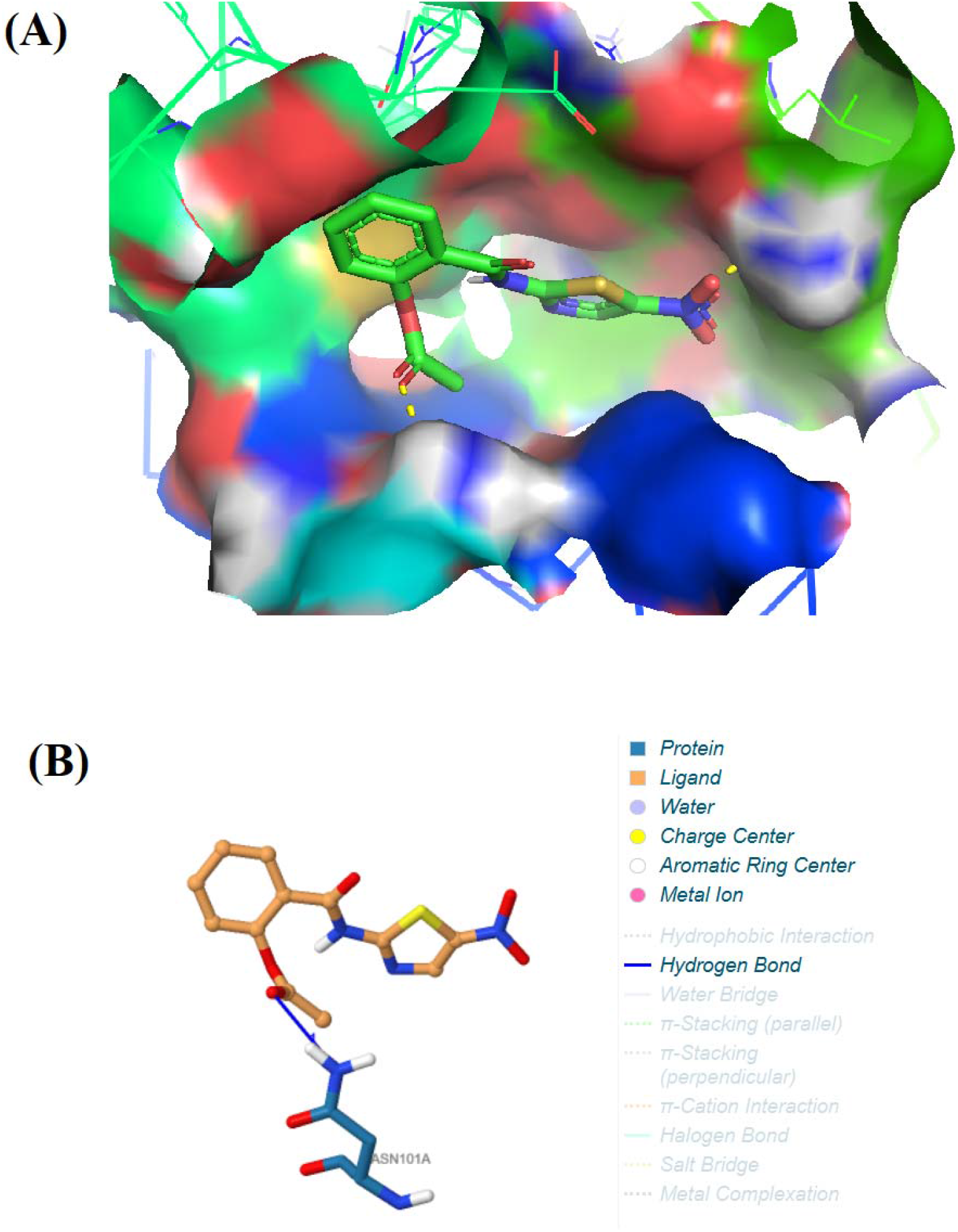
**(A.)** Enlarged view of cavity representation of pocket of SKSR protein of *Cryptosporidium parvum* where drug Nitazoxanide binds; **(B.)** Nitazoxanide-SKSR protein interaction

**Fig. 6:**
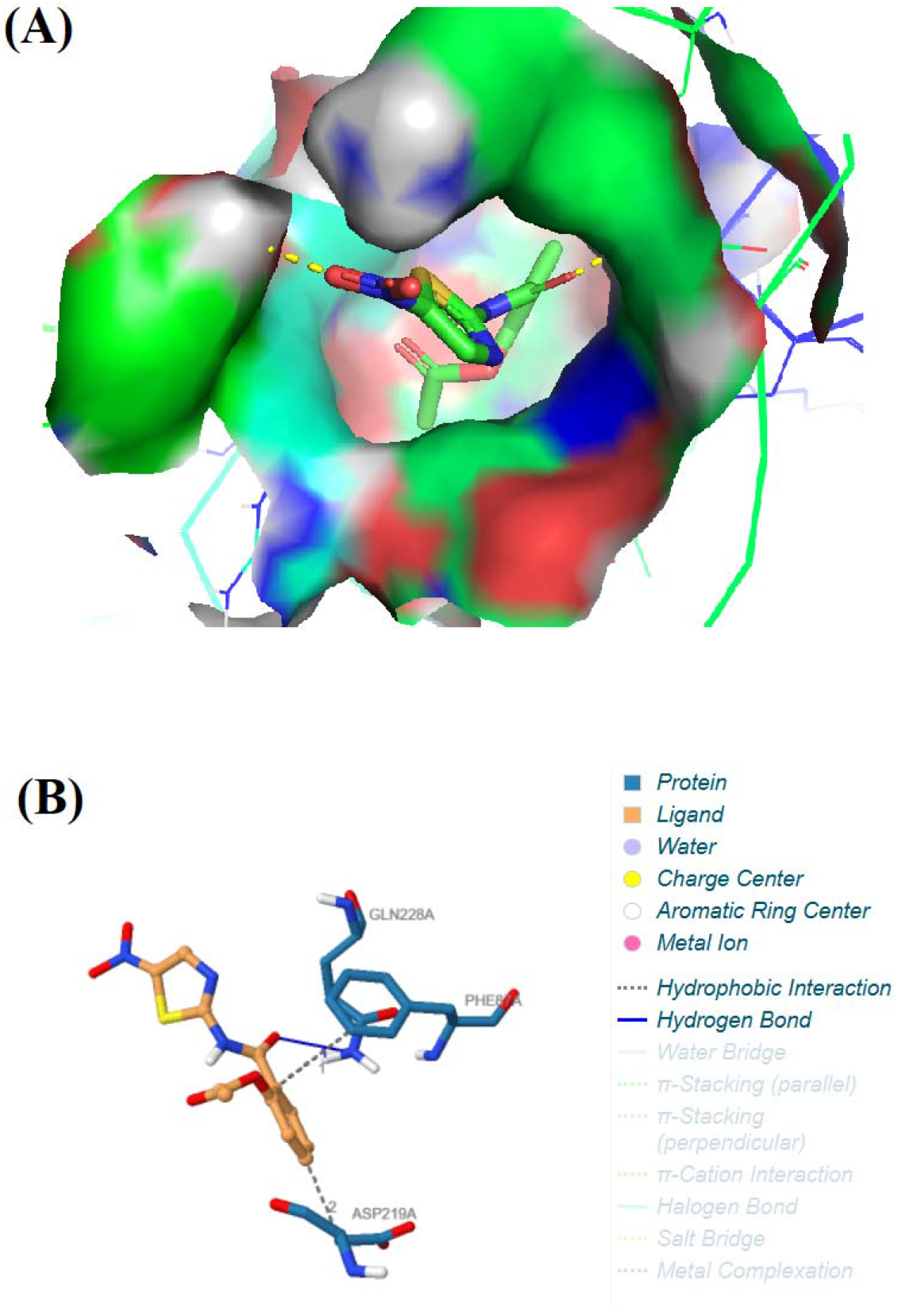
**(A.)** Enlarged view of cavity representation of pocket of key target CpCDPK1 of *Cryptosporidium parvum* where drug Nitazoxanide binds; **(B.)** Nitazoxanide-CpCDPK1 protein interaction.

### 3.3 Molecular Dynamics Simulations

The stability of the proteins upon binding to the ligands was assessed by performing all-atom MD simulations for 100ns using the GROMACS package. Root mean square deviation (RMSD) for the docked complexes along with the apo-proteins was calculated using the gmx rms tool. No major variations were observed upon binding of Artemisinin and Mebendazole to SKSR (Figure 7A, Table 5). A slight fluctuation was observed upon binding of Artemisinin to CpCDPK1, whereas the mean RMSD value for CpCDPK1-mebendazole complex is nearly equal to unbound CpCDPK1 (Figure 8A, Table 5). This indicates that the structures of SKSR and CpCDPK1 remain stable upon binding to Artemisinin and Mebendazole. The variations at the residue level of the protein structure were measured using root mean square fluctuation (RMSF). Minor fluctuations in the average RMSF values are observed between SKSR, CpCDPK1, and their respective complexes (Table 3). Thus, it could be inferred that the protein structures remain stable upon binding of Artemisinin and Mebendazole (Figures 7B, 8B). The radius of gyration (Rg) is a measure to assess the compactness of the protein structure. The average Rg values of CpCDPK1 and its complexes are comparable (Figure 7C). The binding of Artemisinin to SKSR slightly enhances the compactness of the protein, thus making it slightly more stable as compared to Mebendazole (Table 5, Figure 8C). The findings imply that both proteins maintain their stability upon binding of Artemisinin and Mebendazole. Solvent accessible surface area (SASA) refers to the surface area of the protein exposed to the surrounding solvent molecules. Thus, it is used to assess the extent of unfolding of the protein. Minor deviations in the average SASA values were observed over 100 ns of simulations (Table 5, Figures 7D & 8D). Hence, it could be inferred that the proteins retained their respective conformations and remained stable during the course of MD simulations.

**Table 3:** Molecular docking studies of Artemisinin with key targets SKSR protein and CpCDPK1 of *Cryptosporidium parvum*.

| S.No. | Ligand docked with receptor complex | Binding energy (kcal/mol) | Interacting amino acids |
| --- | --- | --- | --- |
| 1. | Artemisinin (Ligand) + SKSR protein (Drug Target) | -6.72 | 58A TYR, 59A ILE, 61A ILE, 86A GLU, 87A LYS, 88A VAL, 92A ASN, 111A ARG, 60A GLU, 89A HIS, 91A THR, 92A ASN, 193A GLN, 288A ARG, 307A LYS, 309A ARG, 209A PHE |
| 2. | Artemisinin (Ligand) + CpCDPK1 (Drug Target) | -9.18 | 100A GLN, 102A TYR, 378A THR, 154A LEU, 379A LEU |

**Table 4:** Molecular docking studies of Nitazoxanide with key targets SKSR protein and CpCDPK1 of *Cryptosporidium parvum*.

| S.No. | Ligand docked with receptor complex | Binding energy (kcal/mol) | Interacting amino acids |
| --- | --- | --- | --- |
| 1. | Nitazoxanide (Ligand) + SKSR protein (Drug Target) | -4.2 | 101A ASN |
| 2. | Nitazoxanide (Ligand) + CpCDPK1 (Drug Target) | -4.81 | 87A PHE, 219A ASP, 228A GLN |

**Table 5:** Parameters calculated for unbound SKSR and CpCDPK1 along with their complexes after 100 ns of MD simulations.

| **System** | **Average**<br>**RMSD**<br>**(nm)** | **Average**<br>**RMSF**<br>**(nm)** | **Average**<br>**Rg (nm)** | **Average**<br>**SASA (nm<sup>2</sup>)** |
| --- | --- | --- | --- | --- |
| **CpCDPK1** | 0.276961 | 0.136363 | 2.48795 | 235.403 |
| CpCDPK1-Artemisinin | 0.343922 | 0.165593 | 2.51608 | 240.786 |
| CpCDPK1-Mebendazole | 0.277603 | 0.143676 | 2.45978 | 234.552 |
| **SKSR** | 3.17718 | 1.27437 | 4.84541 | 695.411 |
| SKSR-Artemisinin | 2.86336 | 0.81046 | 4.58786 | 657.796 |
| SKSR- Mebendazole | 2.62413 | 0.932559 | 5.14367 | 670.277 |

**Fig. 7:**
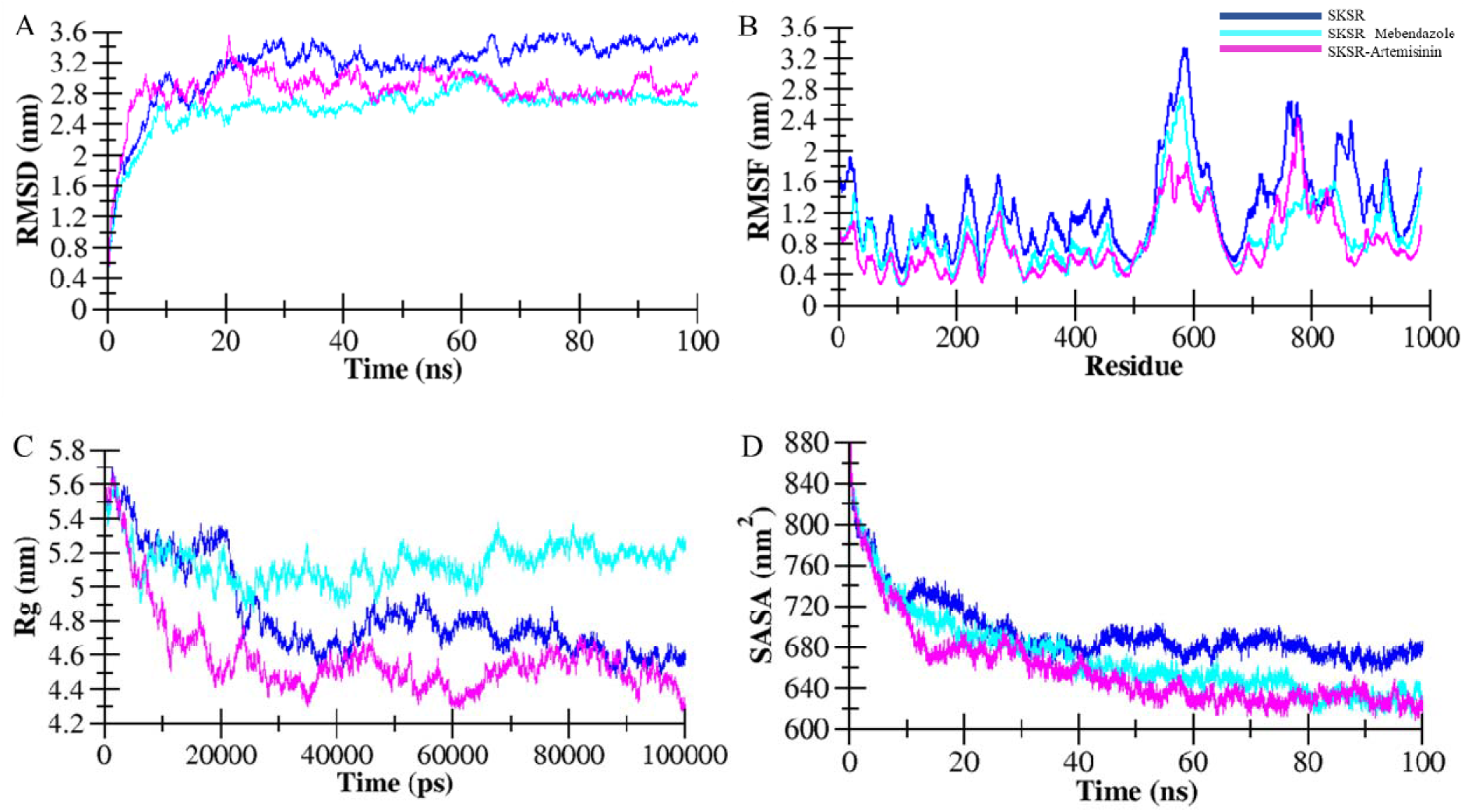
Structural dynamics of SKSR upon binding to Artemisinin and Mebendazole, **(A)** RMSD plot, **(B)** RMSF plot, **(C)** Rg plot, and **(D)** SASA plot

**Fig. 8:**
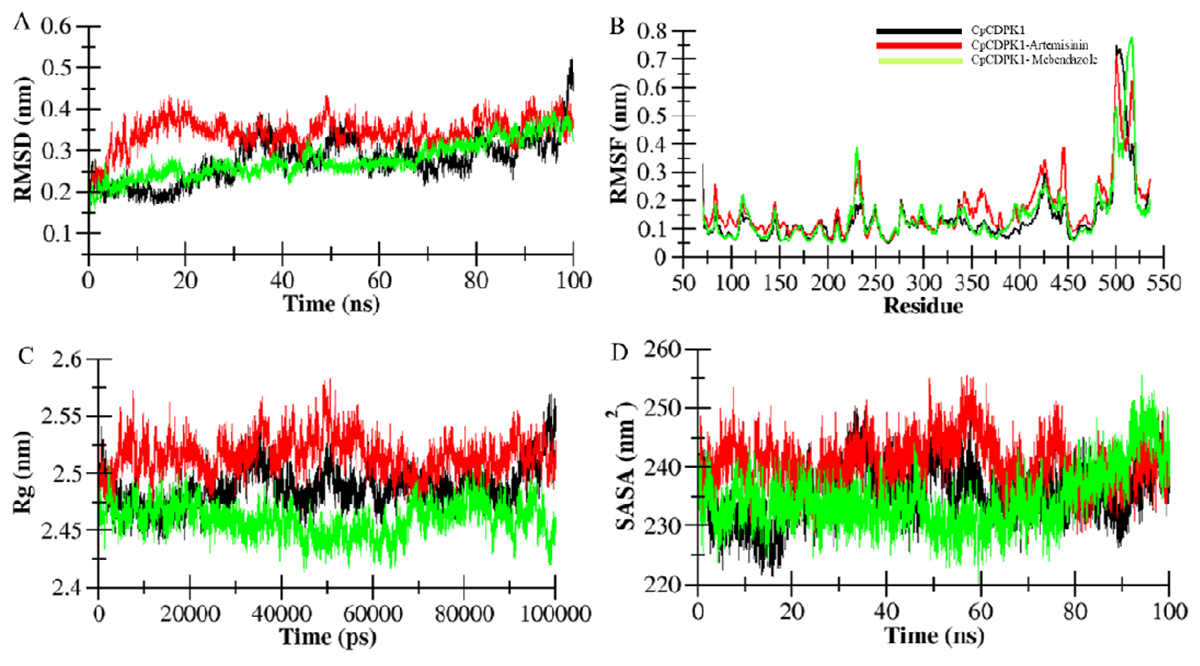
Structural dynamics of CpCDPK1 upon binding to Artemisinin and Mebendazole, **(A)** RMSD plot, **(B)** RMSF plot, **(C)** Rg plot, and **(D)** SASA plot

## 4. Conclusion

The *in silico*-based comparative evaluation of the two repurposed drugs clearly indicates that Artemisinin shows a more pronounced inhibitory potential against *Cryptosporidium parvum* than Mebendazole. This is primarily supported by its consistently lower binding energies across both targets, SKSR and CpCDPK1, suggesting better and more favourable protein-ligand interactions. Notably, the higher affinity of Artemisinin toward CpCDPK1, a key regulator of parasite motility, invasion, and intracellular survival, underscores its potential to disrupt essential signaling pathways critical for parasite viability. We compared the docking results of the above drugs with the current in use recommended drug, Nitazoxanide, and found that the docking results of this recommended drug were very poor with both receptors.

Along with molecular docking, MD simulation further supports the observations by demonstrating that Artemisinin-bound complexes maintain structural stability with reduced fluctuations and improved compactness, particularly in the case of the SKSR protein. This kind of stability indicates the formation of energetically favourable and conformationally stable interactions, which are important for the effective target inhibition. In contrast, although Mebendazole also exhibits stable binding, its comparatively weaker interaction energies and marginally higher structural deviations suggest a less efficient inhibitory profile.

This dual-target engagement of Artemisinin enhances its therapeutic relevance, as simultaneous disruption of virulence-associated proteins (SKSR) and essential kinases (CpCDPK1) may lead to a more comprehensive suppression of parasite growth and pathogenicity. These findings collectively position Artemisinin as a more promising candidate for repurposing against cryptosporidiosis, while Mebendazole may serve as a supportive or secondary option. However, *in vitro* studies are required to further validate these *in silico* findings.

## Acknowledgements

The authors are thankful to the Department of Zoology, University of Allahabad, Uttar Pradesh, India; Department of Zoology, Rajiv Gandhi University, Arunachal Pradesh, India, and Department of Biosciences, Jamia Millia Islamia, New Delhi, India.

